# Age deceleration and reversal gene patterns in dauer diapause

**DOI:** 10.1101/2025.04.14.648662

**Authors:** Khrystyna Totska, João C. V. V. Barata, Walter Sandt, David H. Meyer, Björn Schumacher

## Abstract

The aging process is characterized by a general decrease in physical functionality and poses the biggest risk factor for a variety of diseases such as cancer, cardiovascular diseases, and neurodegenerative disorders among others. Understanding the naturally evolved mechanisms that slow aging and rejuvenate an animal could reveal important concepts how to prevent age-associated diseases and even revert aging. The *C. elegans* dauer state is a robust and long-lived alternative developmental state that after dauer exit has a normal adult lifespan with fully retained fecundity. To understand how longevity during dauer and rejuvenation following dauer exit is mediated, we characterized the gene expression changes during dauer and upon exit. We assessed how biological age, as determined via BiT Age, a transcriptome aging clock, is affected during dauer and upon dauer exit. During the dauer stage, we measured a decelerated increase in age compared to the chronological age and an age reversal following dauer exit. Transcriptomic analyses revealed major metabolic shifts and enhanced biomolecular degradation that are reversed during exit. Moreover, we show that transcription-blocking lesions can induce lasting transcription stress in dauers that is rapidly resolved by transcription-coupled nucleotide excision repair during dauer exit. Our data provide new insights into the underlying mechanisms of naturally occurring age deceleration and rejuvenation.

## Introduction

The aging process progresses throughout life and increases the risk of age-associated diseases and mortality. Some species display an extraordinary degree of plasticity in the rate of aging and, consequently, lifespan. The nematode *Caenorhabditis elegans* has become an important model in aging research in part due to the large degree of lifespan plasticity that is genetically controlled by conserved mechanisms such as the insulin-like growth factor-1 receptor homolog DAF-2^1^. Under adverse conditions such as overcrowding, food scarcity, or heat stress, *C. elegans* can enter the so-called dauer diapause state during its development^2^. This developmentally regulated process serves as a survival strategy amid unfavorable conditions, during which metabolism is typically repressed^3^. The nematode has several diapause states, of which the dauer state is the most robust and long-lived one^2, 4^. In *C. elegans* most cell divisions occur during early embryonic development and already in the earliest larval stage most somatic cells differentiate and only expand in size during subsequent developmental growth. In dauers, the somatic cells are entirely postmitotic thus requiring rejuvenation of terminally differentiated cells, while the arrested germ cells regenerate the germline when resuming to adulthood^5^. Age- and senescence-related molecular changes acquired during diapause states are to a large extent reverted upon diapause exit^3, 6^. While the reports about post-dauer fitness vary, adult lifespan does not decrease in comparison to uninterruptedly developing animals^7, 8, 9^. However, long dauer duration decreases the chances of successfully resuming development, with brood size being the most varied characteristic^7, 8, 9, 10^.

In the past decade, aging clocks have been developed to measure the biological age of organs, organisms, or even entire taxa^11, 12^. Most aging clocks rely on epigenetic marks, particularly age-dependent CpG methylation changes. However, *C. elegans* is devoid of methylated CpG sites. Transcriptomic data reflect a convergence of epigenetic modifications and chromatin structure changes, making it a useful readout for building predictors of age-related changes. Therefore, a transcriptome-based aging clock such as BiT Age can provide highly accurate biological age prediction in *C. elegans*^13^.

In this study, we aimed to uncover the longevity and rejuvenation mechanisms that *C. elegans* employs during dauer diapause and upon its exit. We first performed experiments to ascertain post-dauer fitness in our experimental setup. We observed that nematodes which underwent dauer do not show developmental delays, reduced lifespan, or changes in brood size, regardless of dauer duration. By applying the BiT Age transcriptome-based aging clock, we predict a slowdown of the aging rate during dauer diapause and an age reversal upon dauer exit. We dissect the gene expression programs during the aging process in dauer that is devoid of phenotypic age-related degeneration and then characterize the transcriptome basis for the rejuvenation program occurring during the dauer exit. We determine shifts in metabolic processes and proteostasis mechanisms during diapause that are reversed upon exit. We then show that DNA damage can induce lasting transcription stress as assessed by the gene length-dependent transcription decline (GTLD) that has been observed during aging in multiple species. During the dauer exit transcription-coupled nucleotide excision repair (TC-NER) rapidly restores the transcription integrity indicative of restored genome stability. Our data provide insight into slowed diapause aging free from pathology and a naturally occurring rejuvenation process that could guide geroprotective and rejuvenating intervention strategies.

## Results

### Influence of dauer arrest on post-dauer health

Since reports about consequences of dauer arrest on post-dauer health and fitness are varied, we first performed lifespan, brood size, and developmental assays to assess the impact of dauer diapause on post-dauer fitness in our experimental setup. We chose to use a dauer-constitutive (Daf-c) mutant strain of *C. elegans daf-2(e1370)* to achieve dauer entry with high efficiency and synchronization. We induced dauer entry by incubating synchronized populations of *daf-2(e1370)* L1 larvae at 25 °C (**Fig. 1A**). After 72 hours at 25 °C, all nematodes established dauer arrest, and we termed them day 1 dauers (D1). Dauer recovery was induced by transferring the animals to plates that were freshly seeded with *E. coli* as food source and shifting the incubation temperature to 15 °C.

**Figure 1.**
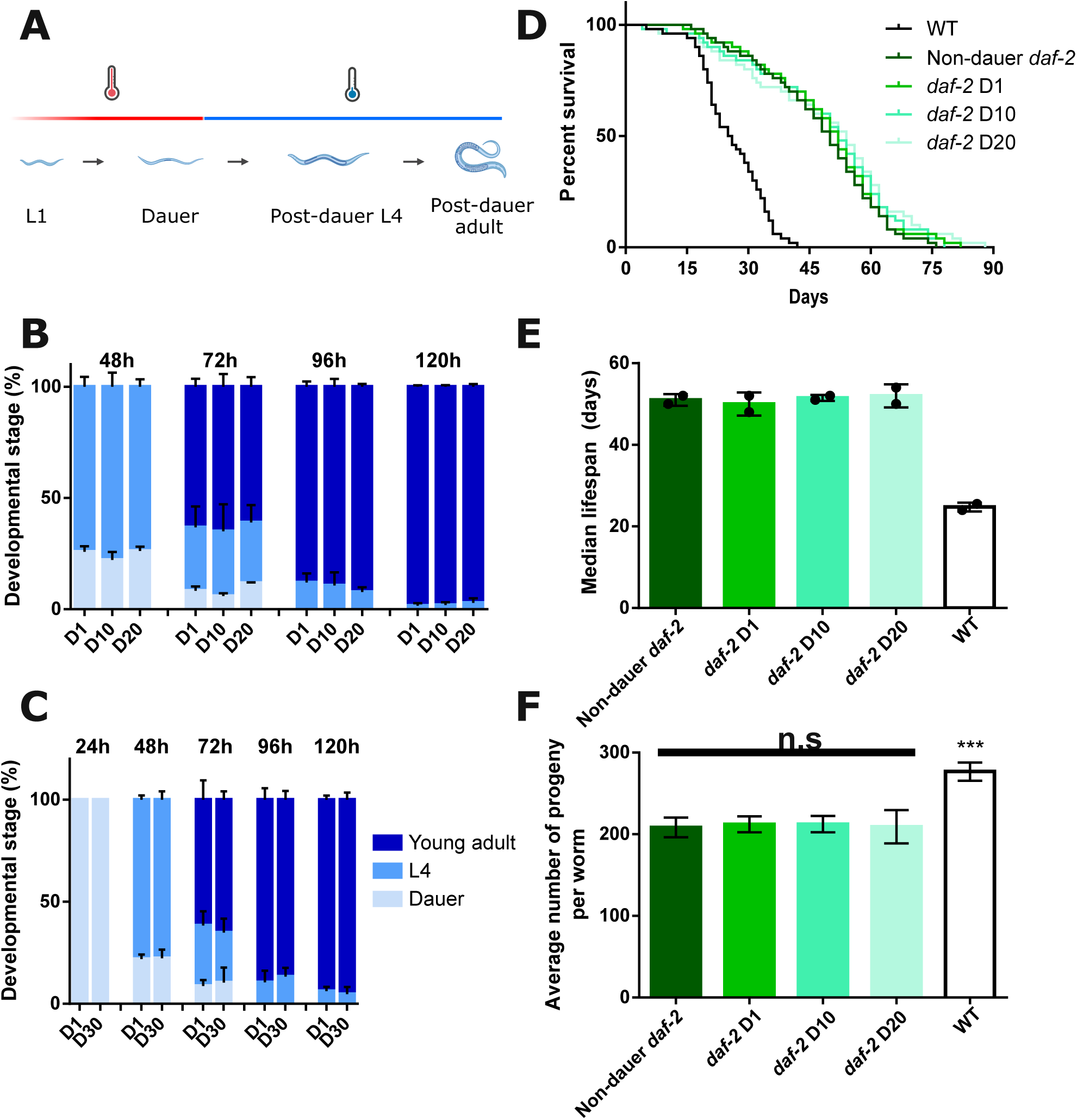
Increasing duration of dauer arrest does not affect post-dauer health and fecundity. **(A)** Schematic overview of the dauer entry protocol. L1 larvae were incubated at 25 °C (red) to induce dauer entry. Dauer exit was induced by shifting the temperature to 15 °C (blue). **(B, C)** Developmental resumption assay of *daf-2* animals arrested in dauer for 1, 10 or 20 days **(B)** and 1 or 30 days **(C)**. At each indicated timepoint after diapause exit induction, the percentage of animals in dauer (grey), L4 (light blue), or young adult (dark blue) stage is shown. **(D, E)** Lifespan assay of wild-type (WT) and *daf-2* animals that did not arrest in dauer, and *daf-2* animals arrested in dauer for 1, 10, or 20 days. **(F)** Brood size assay of wild-type (WT) and *daf-2* animals that did not arrest in dauer, and *daf-2* animals arrested in dauer for 1, 10, or 20 days. One representative experiment is shown in **(B),** and **(C)** shows the median ± SD of two independent experiments per condition. In **(D, E, F)**, the summary of 3 independent experiments is shown as mean ± SD. (A) Mantel-Cox Log Rank test was used for **(B)** (comparisons with WT are shown), one-way ANOVA with Dunnet’s multiple comparison test was used for **(F)**, (comparisons with WT are shown); *** = p<0.001.

We performed a developmental assay in which we assessed the dynamics of dauer recovery of populations arrested in dauer for 1, 10, 20, or 30 days. We observed no delay in populations maintained in dauer for longer periods, with most nematodes reaching the young adult stage 96 h after dauer recovery was induced (**Fig. 1B, C, statistics in Suppl. Table 1**). We also evaluated post-dauer lifespan and brood size in wild-type (WT) and in *daf-2* mutant animals that either did not arrest in dauer (kept at 15°C during development) or arrested in dauer for 1, 10, or 20 days (**Fig. 1D-F**). As reported previously^1^, *daf-2* mutants displayed an extended lifespan (**Fig. 1D,E**) but reduced brood size when compared with WT (**Fig. 1F**). Importantly, we detected no difference in either lifespan or brood size between populations arrested in dauer for different durations and non-dauer *daf-2* worms.

Overall, these results demonstrate that no lasting functional decline occurs in post-dauer nematodes, despite their higher chronological age compared to those that never underwent dauer arrest. These results align with previous reports indicating that although certain age-related molecular changes emerge during diapause, they are largely reversed upon recovery^3, 6^, resulting in sustained post-dauer health and reproductive capacity.

### Predicted biological age during dauer arrest and upon recovery

The dauer diapause exemplifies lifespan plasticity: even though they can outlive normally developing animals several times, once recovered from dauer, they show a normal adult lifespan despite their advanced chronological age (**Fig. 1**). This, together with previous reports on molecular changes during diapause aging in *C. elegans*^2, 3, 6^, suggests that aging is slowed during diapause and reversed during exit.

To investigate this hypothesis, we devised an experimental setup (**Fig. 2A**) in which we collected *C. elegans* populations arrested in dauer for 1, 4, 15, or 30 days, and populations undergoing dauer exit (6 h and 24 h post exit induction), and performed bulk RNA-sequencing. Since the nematodes are still actively recovering from dauer at 6 and 24 h post-exit (**Fig. 1B, C**), we hypothesized that the regulatory transcription programs would be actively engaged during these early recovery phases. We additionally included L3 larvae samples as a non-dauer control group.

**Figure 2.**
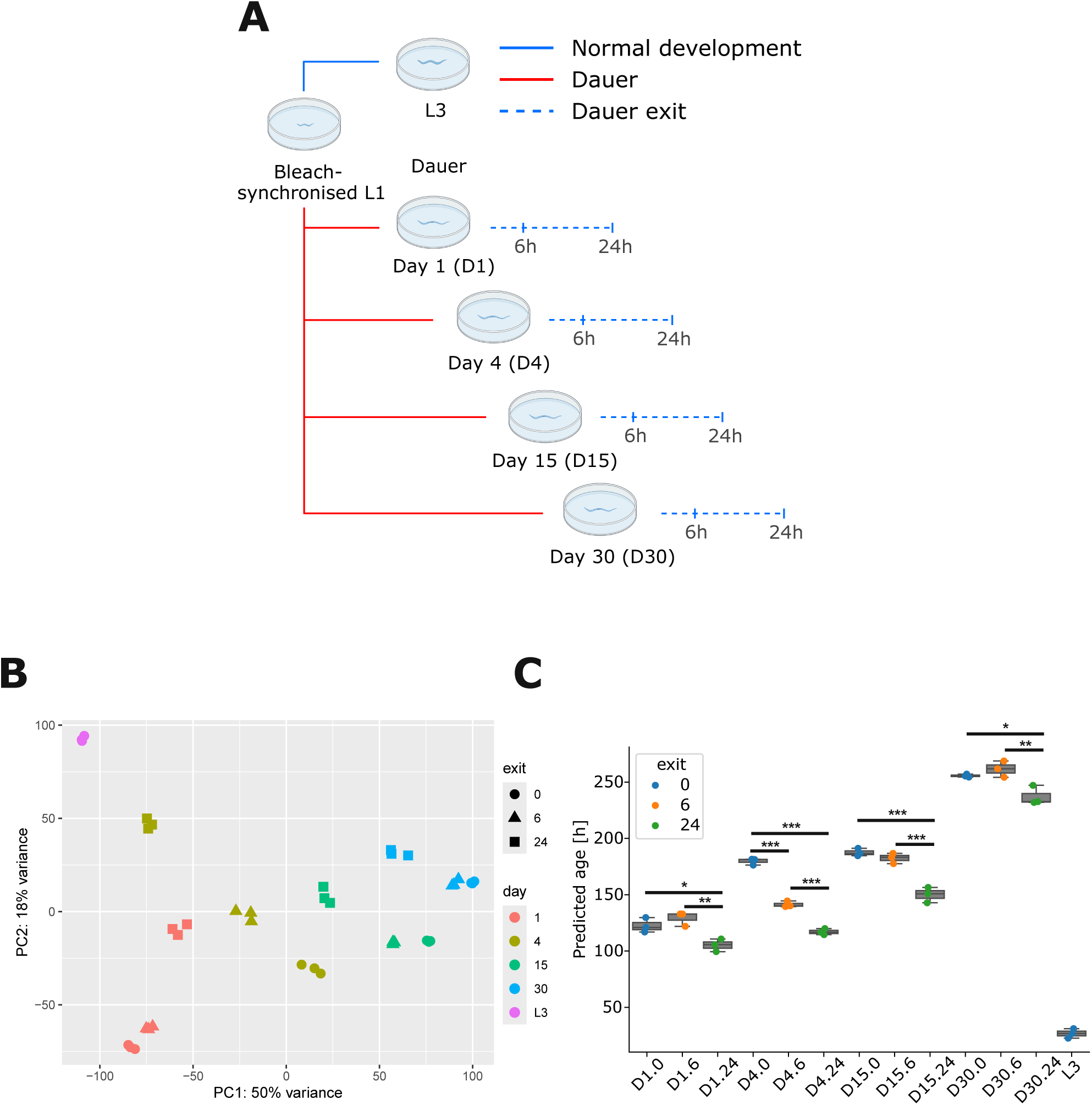
Predicted biological age increases with duration of dauer arrest but is reverted upon dauer exit. **(A)** A schematic representation of the experimental setup for transcriptomic analysis. Blue and red lines indicate plate incubation at 15°C or 25°C, respectively. **(B)** Principal component analysis (PCA) of bulk RNA-sequencing results grouped by developmental stage (color) and time after dauer exit (shape). **(C)** Biological age prediction for dauer and L3 samples using the BiT Age transcriptomics clock. Each dot represents a single RNA-seq sample. The x-axis nomenclature features first the respective day, followed by the number of hours post-exit after the dot. One-way ANOVA with post hoc Tukey test for every day. (p≤0.05 *, p≤0.01 **, p≤0.001 ***)

We first performed principal component analysis (PCA) to identify major sources of variation across samples and visualize sample clustering in lower-dimensional space (**Fig. 2B**). Dauer-arrested samples (called “exit 0”) largely aligned along PC1 in the order of their respective day, indicating progressive changes during the dauer state. Upon dauer exit, most samples shifted in the direction of the L3 control samples. This shift was most pronounced in the D4 dauer exit samples, where both 6 h and 24 h post-exit timepoints showed the greatest proximity to L3. D15 and D30 exit samples also moved toward L3 but to a lesser extent, indicating a less complete transcriptomic recovery trajectory compared to the exit from D4. In contrast, D1 dauer exit samples showed a distinct trajectory, moving closer to L3 along PC2 but slightly diverging along PC1. Given that PC1 appears to reflect chronological aging dependent transcriptomic changes, and PC2 developmental differences, this suggests that D1 dauer larvae undergo transcriptomic recovery primarily through developmental transitions. These results suggest that dauer duration influences the rate of transcriptomic recovery, with shorter duration enabling a more rapid transcriptomic recovery, consistent with previous observations of L1 diapause exit^14^. The PCA thus indicates a progressive dauer aging trajectory and a rejuvenation trajectory that is dependent on the duration of the diapause.

Next, we estimated the biological age of *C. elegans* during dauer arrest and upon recovery using the Binarized Transcriptome (BiT) Age clock that we previously established^13^ (**Fig. 2C, statistics in Suppl. Table 2**). BiT Age uses a binarization approach by converting expression values into binary states (0 or 1) depending on whether the value exceeds the median expression value within each sample. This binarization reduces the variation in gene expression levels that hampered the accuracy of previous transcriptome clocks. Moreover, the BiT Age clock is trained to determine biological age as the lifespan data were known for all training datasets and thus allowed the temporal rescaling of chronological to biological age (i.e. age relative to the time of median survival). We previously showed that the BiT Age clock provides highly accurate predictions of lifespan already during early adulthood and thus reflects the biological age.

Although the BiT Age clock was trained on transcriptomic data from adult *C. elegans*, it indicated a predicted biological age (PBA) progression of chronologically aging dauer-arrested animals. The PBA increased from D1 to D30, with a temporary plateau between D4 and D15, suggesting a non-linear aging trajectory. The rate of biological aging progressively slowed over time: for example, at D1, the PBA is 123 h, while at D4 it was 180 h. This corresponds to a difference in biological age of 57 h over a chronological time span of 72 h. Therefore, the biological aging rate between these two timepoints is ca. 0.79 (57 h/72 h). Since, at D30, the PBA was 256 h, the rate of biological aging is ca. 0.12 (76 h/624 h) between D4 and D30. These results indicate that although biological aging continues during the dauer state, the rate of aging decreases sharply over time.

Additionally, the PBA at 24 h post exit at D15 is younger than at 24 h post exit at D30 suggesting that older dauers require more time to reverse age-associated changes accumulated during diapause. However, despite this delay in transcriptomic recovery, the results of our developmental assay (**Fig 1. D,E**) indicate that older dauers do not suffer phenotypic consequences in terms of the developmental dauer exit program. This suggests that longer diapause duration delays molecular recovery but does not impair the developmental control of dauer recovery.

In general, the PCA results align well with BiT Age predictions. The chronologically and biologically youngest D1 dauers showed weak rejuvenation upon exit according to BiT Age, consistent with their limited shift along PC1, while D4 exit samples showed the strongest dauer exit dynamics according to PCA, which is mirrored by the largest reduction in predicted biological age. Taken together, these results demonstrate that the biological age progression slows during dauer and is reversed upon dauer exit, with a pace of recovery depending on the duration of the dauer diapause.

### Transcriptomic patterns throughout dauer arrest and during recovery

We next sought to characterize the gene expression dynamics during dauer aging and exit. Instead of using multiple pairwise comparisons, we used a statistical approach that models the trajectory of each gene’s expression over the chronological time. This enabled us to assess whether the expression of a gene increases or decreases over sequential timepoints, thereby revealing broader regulatory trends. We focused on (1) dauer aging: changes in gene expression from D1 to D30 of the dauer stage, and (2) dauer exit: changes in gene expression from the first 6 to 24 hours after induction of dauer exit. For each gene, we used a linear model to compute a slope that quantifies both the direction (up- or downregulation) and rate of expression change over time across dauer aging and dauer exit (see Methods for details). Genes with positive slopes increased in expression over the time course, whereas those with a negative slope decreased. We then identified differentially expressed genes (DEGs) as those with a statistically significant slope, e.g. a consistent increase in expression across the time course and performed a KEGG pathway enrichment analysis to detect patterns in biological pathways.

### Dauer aging

During dauer aging, 24 pathways are significantly downregulated, whereas only 5 pathways are significantly upregulated. Multiple metabolic pathways are downregulated (**Fig. 3A, Suppl. Fig. 1, Suppl. Table 3**), consistent with the low-metabolic, stress-resistant nature of the dauer state. Given the lifespan extending effects of dampened metabolic activity, this metabolic downregulation aligns with the BiT Age prediction that the biological aging rate during the dauer state decreases. Conversely, longevity regulating pathways, proteasome, and autophagy are significantly upregulated during dauer aging. This suggests that dauer larvae might engage enhanced protein quality control and cellular maintenance mechanisms that help mitigate damage accumulation and potentially contribute to their extended survival.

**Figure 3.**
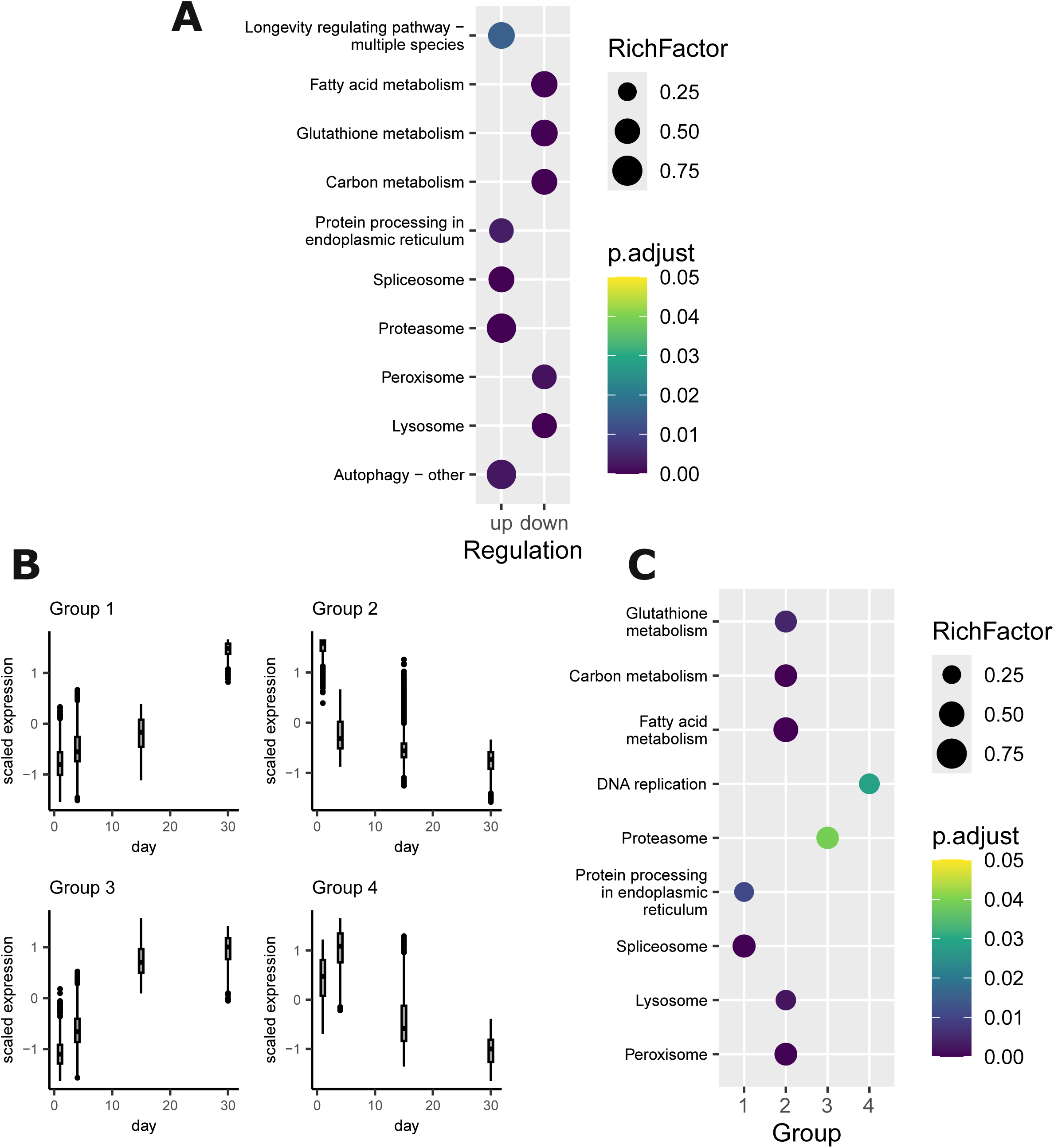
Dauer diapause transcriptomes reveal dauer longevity-associated pathways. **(A)** A subset of KEGG pathways enriched for genes exhibiting differential expression over the dauer aging time course. Color-coded are the adjusted p-values, and the bubble size represents the rich factor. **(B)** Genes were grouped into four clusters based on their expression patterns over the dauer aging time course (days 1, 4, 15, and 30) using k-means clustering. The y-axis represents scaled expression values, and the x-axis shows the chronological age in days. Cluster sizes: Group 1 (1,826 genes), Group 2 (3,082 genes), Group 3 (1,754 genes), Group 4 (1,544 genes). **(C)** Subset of enriched KEGG pathways for each of the four gene clusters identified across the dauer aging time course. The x-axis represents the gene clusters (Groups 1–4), while the y-axis lists selected pathways. Color-coded are the adjusted p-values, and the bubble size represents the rich factor.

To further characterize expression dynamics beyond linear changes, we clustered DEGs into four groups based on their dauer aging-dependent expression profiles (**Fig. 3B**). While the linear model analysis identified general trends, clustering allowed us to identify distinct co-regulated gene sets that change in non-linear pattern across the dauer time course. The largest cluster (Group 2) showed a clear downregulation and is enriched for metabolic pathways such as the glutathione, carbon, and fatty acid metabolism (**Fig. 3C, Suppl. Fig. 2**). This reflects the metabolic decline observed in the initial pathway analysis (**Fig. 3A**) and further suggests that broad metabolic downregulation is a dominant feature of dauer aging.

Group 3 showed a consistent upregulation and is significantly enriched for the proteasome, indicating a potential role of cellular maintenance in the longevity of the dauer state (**Fig. 3C, Suppl. Fig. 2)**. This suggests that while dauer larvae reduce overall metabolic activity, they simultaneously enhance proteostasis mechanisms, possibly as an adaptive strategy to extend lifespan under energy-restricted conditions. Group 1 is similarly showing an upregulation over time, but remains largely stable from D1 to D15, while showing a sharp upregulation at D30 (**Fig. 3C, Suppl. Fig. 2)**. This group was enriched for spliceosomal and protein processing pathways, similar to the enrichment analysis of the linear slopes (**Fig. 3A**). The clustering approach revealed that this enrichment is primarily driven by the late-stage, D30 dauer, rather than a continuous increase over the chronological age. This suggests that certain transcriptomic responses are uniquely activated in long-term dauer survival (Group 1), potentially as a late adaptation to extended energy deprivation or stress conditions. In conclusion, by integrating the linear model and clustering analysis, the results support a model where dauer larvae dynamically regulate transcriptomic programs depending on their time in dauer, with energy conservation and cellular maintenance continuously regulated, while spliceosomal and protein processing pathways only emerge as a late-stage response.

### Dauer exit

Next, we focused on the characterization of transcriptomic dauer exit dynamics. To evaluate the extent to which the transcriptomic responses to dauer exit overlap across different dauer aging timepoints, we compared the sets of DEGs identified at D1, D4, D15, and D30 using the same linear model approach used earlier. Here, DEGs were defined as genes with a significant change in expression, i.e. a slope significantly different from 0, across the in-dauer, 6 h post-exit and 24 h post-exit samples. Overall, these DEG sets showed a high degree of similarity, indicating the core processes associated with the dauer exit program. D1 exit showed the most distinct exit pattern and displayed the lowest overlap with the other timepoints, while D15 and D30 showed the highest similarity (**Fig. 4A**). These observations are consistent with the exit trajectories observed in the PCA, where D1 samples diverged most significantly from other days.

**Figure 4.**
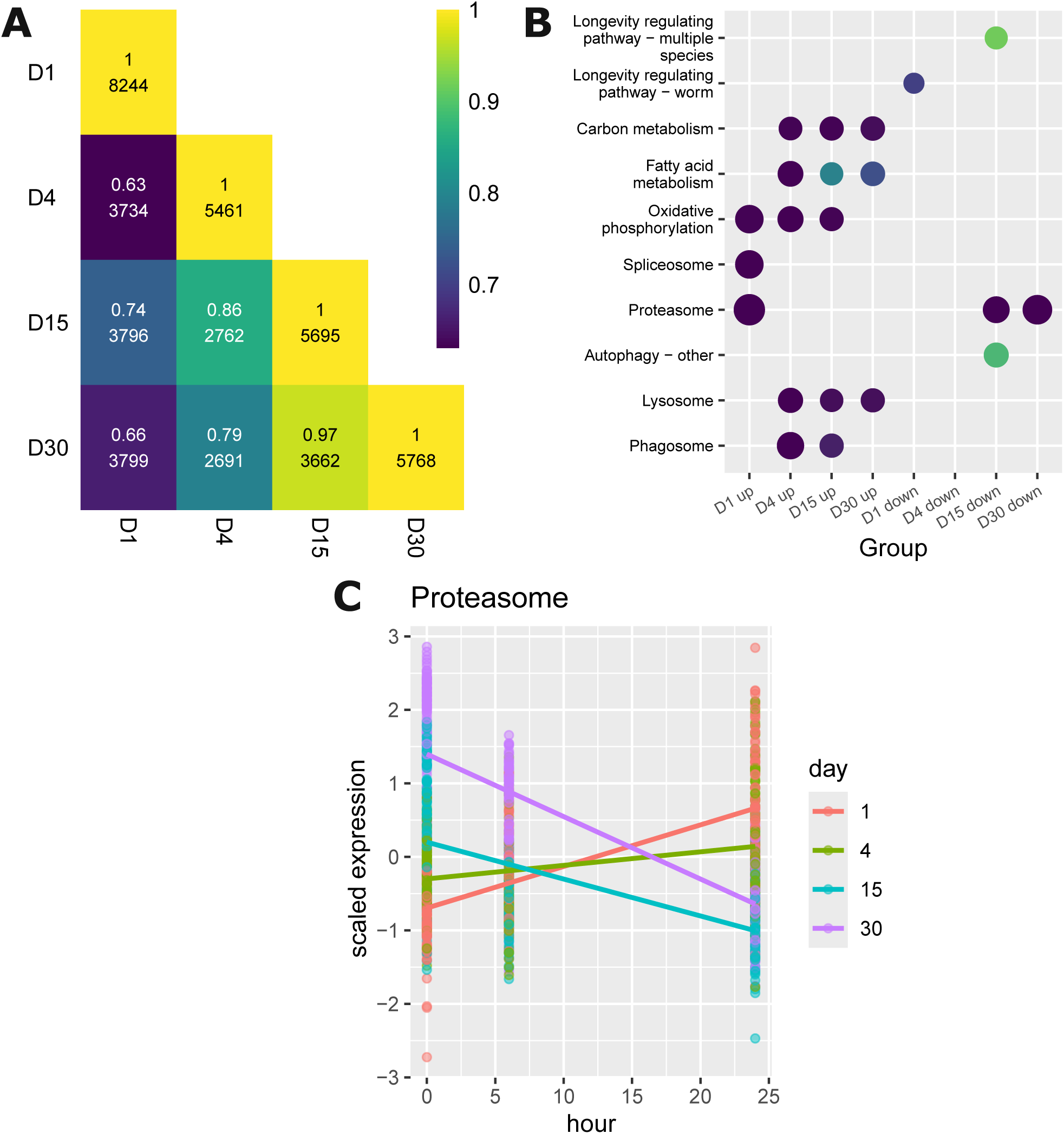
Dauer exit trajectories reveal rejuvenation-associated pathways. **(A)** Heatmap showing the overlap in differentially expressed genes between dauer exit at different chronological ages. The color scale represents the fraction of shared genes that exhibit the same direction of regulation. The upper number in each cell indicates this fraction, while the lower number denotes the total number of overlapping differentially expressed genes between the two time points. **(B)** Subset of enriched KEGG pathways for differentially expressed genes at each dauer exit time point. Color-coded are the adjusted p-values, and the bubble size represents the rich factor. **(C)** Scaled expression values of proteasomal genes are shown across dauer exit time points (0, 6, and 24 hours). Each dot represents the scaled expression of an individual proteasomal gene. Colored lines indicate linear model fits for each aging time point (days 1, 4, 15, and 30), illustrating how proteasomal gene expression exit-dynamics change with dauer aging.

To investigate how the transcriptional response to dauer exit depends on the duration of diapause, we performed KEGG pathway enrichment analysis on differentially expressed genes across dauer exit time points from animals recovered at days 1, 4, 15, and 30 of dauer arrest. The KEGG enrichment analysis showed that dauer exit is generally associated with the upregulation of lysosomal and multiple metabolic pathways (**Fig. 4B**), indicative of a reactivation of energy-demanding processes during dauer exit following the metabolically suppressed dauer state. However, the specific biological pathways enriched during exit vary depending on the dauer duration. This may be a result of cumulative changes to the worms during the dauer state. Furthermore, the enrichment of longevity-associated genes among downregulated ones on D1 and D15 could be interpreted as an ending of dauer-longevity mechanisms and resumption of a normal life cycle.

Among the most strikingly dynamic pathways was the proteasome (**Fig. 4B, C)**. Proteasomal genes were strongly upregulated during dauer aging, suggesting an adaptive upregulation of proteostasis during prolonged diapause (**Figure 3**). Upon dauer exit, however, this trajectory is reversed and proteasomal gene expression showed a distinct time-dependent pattern: proteasomal genes were enriched among upregulated genes during exit from early (D1) dauers, were no longer significantly enriched by D4, and were instead enriched among downregulated genes in animals exiting at later stages (D15, D30) (**Fig. 4B**). At the 24-hour post-exit timepoint, proteasomal expression was highest in D1-exit animals and lowest in D30-exit animals—opposite to the expression pattern seen during dauer aging itself (**Fig. 4C**), suggesting an interaction between the duration of diapause and the ability to dynamically regulate proteasomal activity during recovery (**Fig. 4C**). These data suggest that early-dauer exit, having not yet strongly upregulated proteasome genes during diapause, require an additional transcriptional upregulation upon exit to restore proteostasis. In contrast, long-term dauers already upregulated their proteasomal capacity, enabling a normalization or even downregulation during recovery.

Our findings suggest: (1) that a dampened metabolism and an interplay between protein synthesis and degradation pathways, including lysosome and proteasome activity, is a part of the dauer longevity program, potentially in combination with alternative splicing regulation, and (2) that a transcriptomic program driving rejuvenation during dauer recovery involves proteasomal dynamics and the upregulation of metabolic processes.

### Effects of UV-irradiation on dauer recovery and gene length-dependent transcription patterns

The maintenance of the genome is an essential component of longevity and how genome stability could be reinstated in a rejuvenation process remains unclear. We thus investigated how dauer larvae maintain genome integrity and recover from DNA damage. A particularly striking feature of dauer larvae is their resistance to various types of stress^16^, including UV-induced DNA damage^17^. To explore the ability to maintain their genomes, we used UV as genotoxic paradigm for inducing transcription-blocking DNA lesions, which play an important role, particularly in the aging process of postmitotic cell types such as neurons^18^. We chose UVB treatment because it predominantly induces the well-defined cyclobutane pyrimidine dimers (CPDs). CPDs distort the helix structure resulting in stalling of transcription elongation until repair by transcription-coupled nucleotide excision repair (TC-NER). By interfering with transcription elongation, this lesion type is particularly relevant for non-dividing cell types, such as those comprising the dauers. We therefore exposed worms to UV radiation at the beginning of diapause and assessed recovery dynamics depending on dauer duration.

First, we irradiated D1 worms and induced dauer recovery, either directly or by leaving the irradiated animals until D4 before triggering the exit. With increasing UV dose, the recovery from dauer was delayed suggesting that during the exit process, DNA repair requires time to remove the lesions. However, we observed only a very minor reduction in this delay when the animals were given 4 more dauer days to recover from the UV-induced damage (**Suppl. Fig. 3, statistics in Suppl. Table 7**). This suggests that the repair of UV-induced damage in genes required for dauer exit occurs predominantly after exit is triggered, rather than during the dauer phase itself. To ascertain that despite the absence of a phenotypic consequence in dauers, the applied UV irradiation has a biological consequence and requires the repair of UV-induced DNA damage, we next tested the effects of perturbing the nucleotide excision repair (NER) pathway. NER is the major mechanism required for removing helix-distorting lesions such as UV-induced CPDs. Upon stalling of the RNAPII at the lesions, the CSB-1 protein induces transcription-coupled (TC-) NER. Throughout the genome such lesions are recognized by XPC-1, that initiates global-genome (GG-) NER^19^. Both converge on the NER core pathway by recruiting XPA-1. Upon UV irradiation, mutations in *csb-1* lead to developmental growth delays and premature aging in the worm’s postmitotic somatic cells (as they require transcription but not DNA replication), while *xpc-1* mutants are UV sensitive in proliferative germ cells. *xpa-1* mutants are exquisitely sensitive to UV in all cell types, as no NER activity is present^20^. These nematode phenotypes reflect the human disease syndromes, where TC-NER defects caused by *CSB* mutations lead to growth delays and premature aging, while GG-NER defects caused by *XPC* mutations leads to elevated mutations throughout the genome of proliferating cells resulting in exquisite skin cancer susceptibility^21^.

Given the role of NER in DNA damage-driven aging, we tested the consequences of a perturbed NER pathway for dauer exit by following the dauer recovery of NER mutants (**Fig. 5A-D, statistics in Suppl. Table 5**). All NER mutants had similar recovery dynamics to single *daf-2* mutant in the absence of UV-induced DNA damage (**Fig. 5A**). However, upon UV exposure we observed a dose- and NER-dependent delay in resuming development (**Fig. 5B-D**). Completely NER deficient *daf-2;xpa-1* mutants were unable to resume development even at the lowest dose of 15 mJ/cm^2^. TC-NER deficient *daf-2;csb-1* mutants displayed a transient delay of dauer exit, while at 30 mJ/cm^2^ most TC-NER defective animals were unable to resume development. In contrast, GG-NER deficient *daf-2;xpc-1* mutants showed a similar dauer exit as *daf-2* single mutants. However, in contrast to repair proficient animals, the UV-treated *daf-2;xpc-1* mutants failed to produce offspring consistent with the role of GG-NER to repair UV lesions in germ cells (data not shown). These results are highly consistent with the role of TC-NER in somatic cells, which are mostly postmitotic after embryonic development.

**Figure 5.**
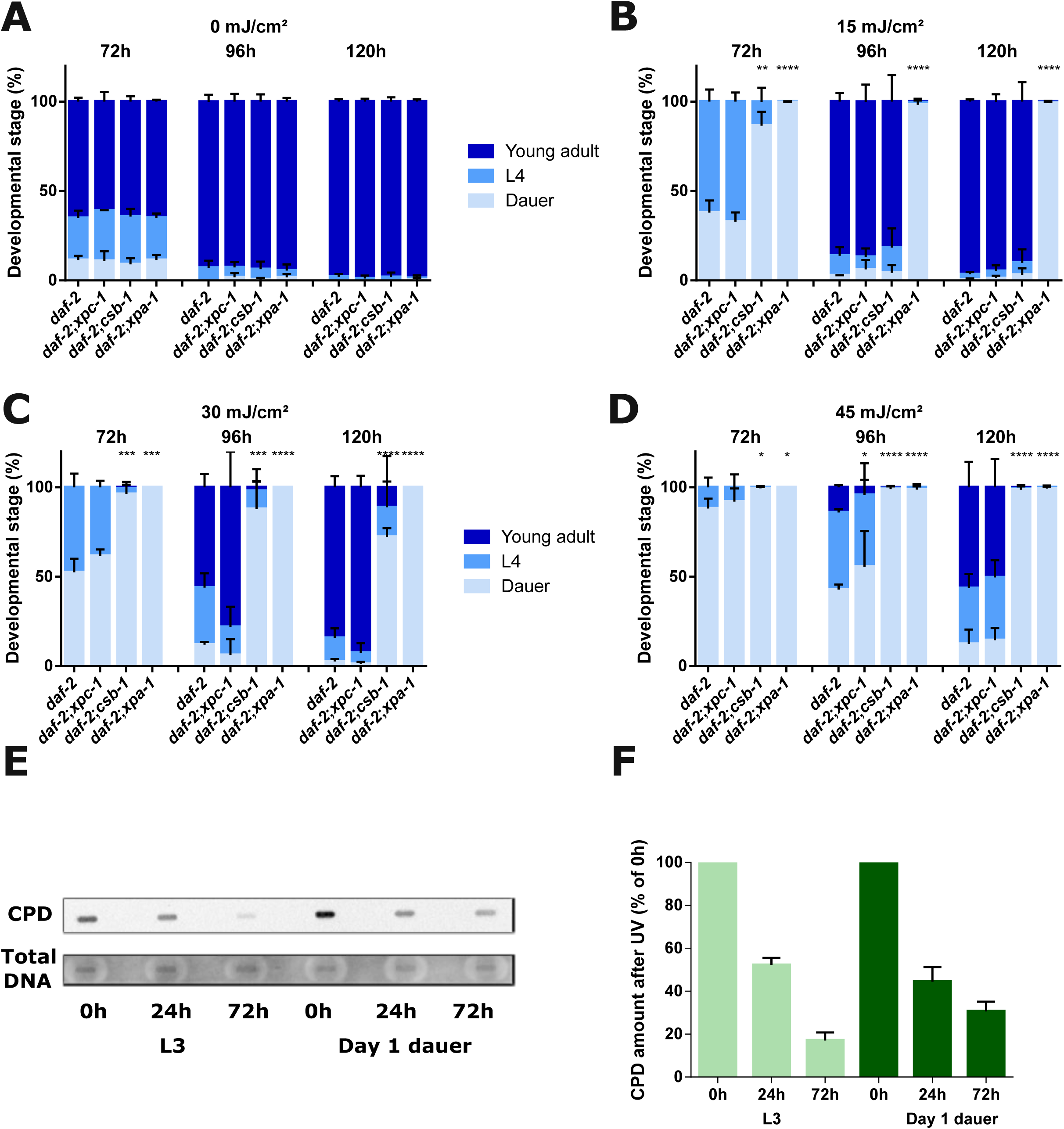
NER machinery is required for successful dauer exit which is impaired by UV treatment equally in D1 and D4 worms. **(A-D)** *daf-2*, *daf-2;xpc-1*, and *daf-2;csb-1, daf-2;xpa-1* animals were UV- or mock-treated at day 1 of dauer. Panels **(A)** to **(D)** show, respectively, 0, 15, 30, and 45 mJ/cm2 conditions. Dauer exit was induced immediately after the UV treatment. (average of n=3 independent experiments per strain and dose is shown, error bars represent the standard deviation (SD); Two-tailed t-test). (E, F) *daf-2* animals were UV-treated (75 mJ/cm2) at the L3 or dauer stage (D1). CPDs were measured via slot blots by antibody staining, 0, 24 and 72h after UV treatment, and normalized to the respective 0h time-points (result shown in panel **(E)** is representative, experiment was repeated 2 times).

To further address the NER activity during dauer, we used an established anti-CPD antibody that specifically recognizes this lesion type to assess the removal of UV-induced CPDs in a slot blot (**Fig. 5 E,F**). The CPD removal kinetics in dauers were almost as efficient in comparison to non-dauer L3 larvae with over 50% of the UV-induced damage repaired in the first 24 h followed by slower repair, in agreement with previous reports that suggest biphasic repair kinetics^22^. The similar recovery dynamics of D1 and D4 dauers despite the additional time the latter had to repair (**Suppl. Fig. 3**) suggests that while global repair mechanisms remain active during dauer, the repair of transcriptionally relevant regions necessary for exit may occur only upon exit initiation. This aligns with the role of TC-NER in repairing actively transcribed regions and may explain why TC-NER-deficient animals exhibit a delayed dauer exit when treated with UV.

While the CPD quantification (**Fig. 5E, F**) assesses the overall burden of DNA lesions throughout the genome, the effect of CPDs on blocking transcription is limited to those occurring on the actively transcribed strand leading to stalling of RNAPII. The effect of transcription blocking DNA lesions can instead be assessed by the reduced expression of genes encoded by long open reading frames. Here, UV lesions lead to the gene length dependent transcription decline (GTLD) indicative of transcription stress. Similarly to the effect of transcription blocking UV-induced lesions, the GTLD is observed during aging in multiple species. It is thought that the GLTD is triggered by the accumulation of transcription blocking lesions during the aging process^23, 24^. Hence, investigating the changes in gene length dependent expression during dauer exit may further elucidate the reversal of the transcription-blocking DNA lesions causing the age-related GLTD. We assessed the gene expression changes in D1-UV-treated animals at D4 and upon dauer exit on day 4 (**Fig. 6, Suppl. Fig. 4A**). In the dauers the downregulated genes in the UV-treated samples relative to the untreated samples are longer than the upregulated genes, indicative of DNA damage-induced GLTD even 4 days after UV treatment (**Fig. 6, Suppl. Fig. 4A**). During dauer recovery (6 h, 24 h), the transcriptomic differences between UV-irradiated and control samples largely normalize (**Suppl. Fig. 4B)**. Only a few genes remain differentially expressed, and those did not show any length differences between up- and downregulated genes, indicating that transcription-coupled repair efficiently restored transcription output and successfully ameliorated the GLTD.

**Figure 6.**
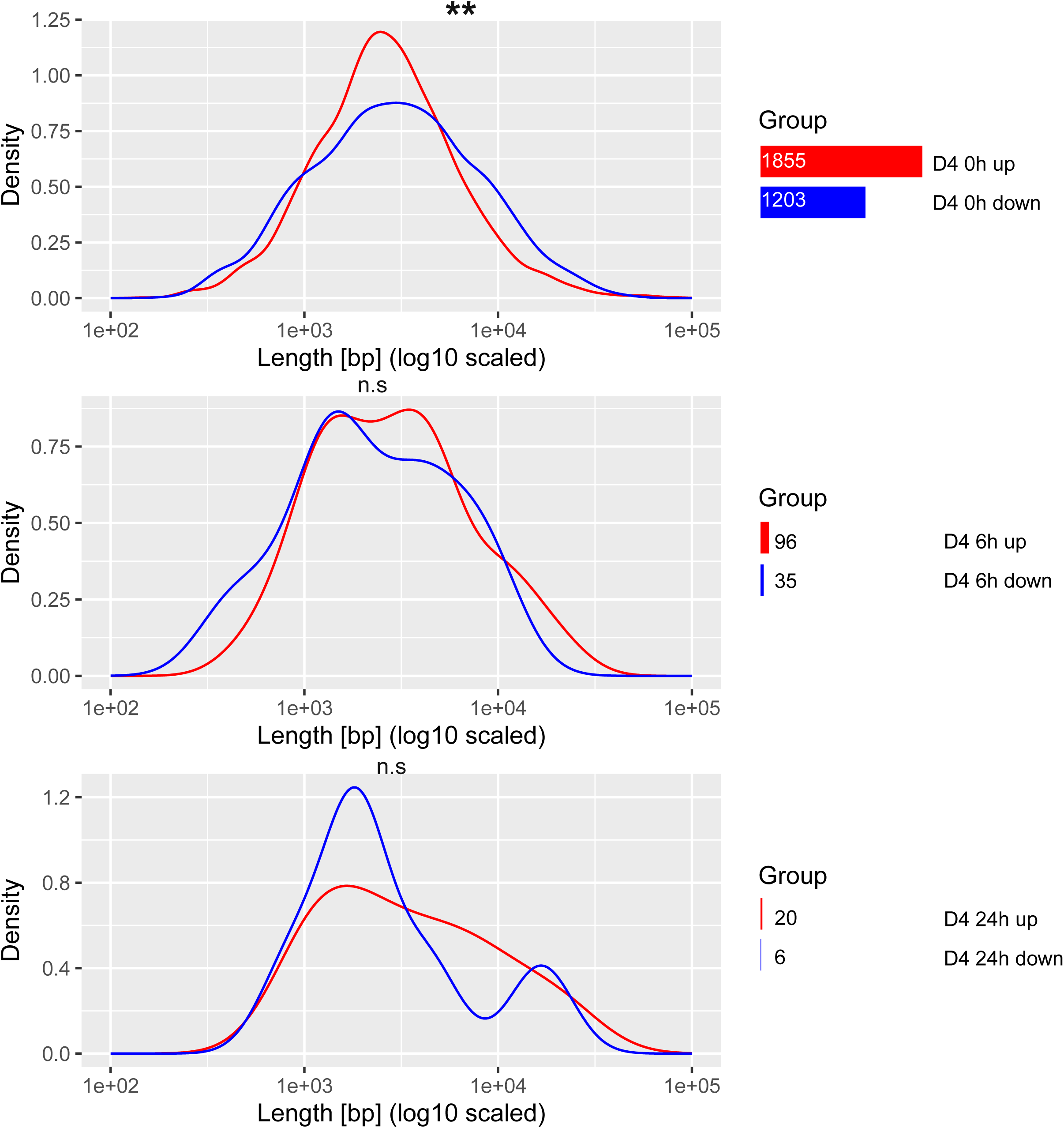
Recovery of DNA damage-induced transcription stress during Dauer exit. Density plots of gene lengths of genes differentially up- or down-regulated in UV-treated samples of dauer exit on D4 relative to untreated samples of dauer exit on D4. Mann Whitney U test was used to identify statistically significant differences in gene lengths between the groups. (p< 0.05 *, p < 0.01 **, p < 0.001 ***). The number of genes that are differentially over- or under-expressed in UV-treated samples of dauer exit on D4 relative to untreated samples of dauer exit on D4 is written to the right and corresponds to the size of the bar.

Overall, these results underscore the remarkable ability of dauer larvae to withstand and recover from genotoxic stress, highlighting the importance of NER, particularly TC-NER, in facilitating successful dauer exit. Additionally, the observed recovery from UV-induced GLTD suggests that the dauer exit process triggers the removal of transcription-blocking lesions and restores transcriptional integrity, enabling efficient reactivation of biological processes upon exit.

## Discussion

The understanding of the underlying mechanisms of naturally occurring slowed aging and rejuvenation could reveal effective paradigms for preserving and regaining the functional integrity of the aging organism. The *C. elegans* diapause is a particularly striking example of lifespan plasticity given that despite a chronological age that can be multiple times longer than a normal lifespan^2^ it recovers to pursue an unchanged adult lifespan with full reproductive capacity. The somatic cells of the dauer are already entirely postmitotic, thus necessitating the preservation of their functional integrity during the diapause stage and rejuvenation without cell renewal upon exit. Here, we employed the *C. elegans* dauer diapause and dauer exit as model for exploring physiological mechanisms of age deceleration and rejuvenation.

We ascertained that in *daf-2* mutants, dauer diapause has no negative consequences on postdauer lifespan, brood size, and development. We produced a bulk RNA-seq dataset that includes multiple timepoints during the diapause and recovery to characterize their transcriptomic patterns. Using the BiT Age transcriptomic clock, we demonstrate that predicted biological age increases during dauer arrest and decreases upon exit induction. The rate of aging changes from 0.8 units of biological time for one unit of chronological time in the first three days to 0.1 from D4 to D30, suggesting an aging plateau is reached during the first days of the diapause arrest. Upon dauer exit, the biological age is reset as recovered dauers show the same lifespan and fecundity as chronologically younger animals that instead underwent a normal larval development. This was reflected by a reversal of the predicted biological age. While this reversal was most pronounced 24 h after 4 days of diapause, more extensive diapause periods result in slower resetting of the predicted age, with negligible changes during the first 6 hours post exit induction, indicating the extended time that is required for the rejuvenation process the longer the dauer diapause lasted.

The *C. elegans* dauer exit model for rejuvenation neither leads to a reentry of post-mitotic differentiated cells into the cell cycle nor a loss of cell identity in stark contrast to Yamanaka reprogramming as a rejuvenation method. Similar to the reversal of the epigenetic clock during reprogramming of somatic cells into induced pluripotent stem cells (iPSCs)^25^, dauer exit reverts the biological age as determined by BiT Age. The longer the dauer diapause lasted, the longer the resetting of the predicted biological age took, indicating a divergence between the processes needed for developmental recovery and those determining biological aging. Eventually, however, the animals are completely rejuvenated as they are set for a normal adult lifespan regardless of the time spent in dauer.

The investigation of enriched biological pathways on a transcriptional level affected by dauer duration and exit can elucidate which mechanisms may be implicated in the reversal of aging during dauer exit and lifespan plasticity during dauer diapause. Specifically, our data indicate that autophagy, the proteasome, and metabolic processes are most prominently regulated during dauer. Autophagy, which was upregulated during dauer aging, plays a crucial role in maintaining regenerative capacity as demonstrated in hematopoietic stem cells in mammals^26^. Autophagy is also induced by dietary restriction and dietary-restriction mimicking drugs, and during OSKM reprogramming^27, 28, 29^. The proteasome is required for the maintenance of proteostasis^30^. Hence, the upregulation of proteasomal activity during dauer aging, paired with a general downregulation of multiple metabolic pathways, could prevent the accumulation and aggregation of damaged proteins and other biomolecules. The general downregulation of metabolic pathways during dauer is very consistent with the lifespan extension triggered by lowered metabolic activity, such as upon dietary restriction. Dysfunctional alternative splicing has recently been associated with aging, as age-related changes in splicing factor levels are detected in mice, rats, and humans ^31, 32^. Moreover, in *C. elegans* some splicing factors are connected to longevity through mTOR and AMPK pathways^31^.

The general upregulation of metabolic activity upon dauer exit suggests a recovery of the dampened metabolic processes during dauer entry and diapause itself. Additionally, the accumulation of aging phenotypes may not proceed in a purely linear manner which is also suggested by the BiT Age results. Different hallmarks of aging have to be ameliorated during dauer exit potentially requiring the activity of different pathways.

Dauers exhibit resilience to UV-induced damage and are proficient in repairing UV-induced lesions during the arrest^17^. Transcription-blocking lesions, which can be experimentally most effectively induced by UV irradiation, play a particularly important role in organismal aging. The accumulation of transcription-blocking lesions was recently shown to lead to transcription stress that results in the age-related GLTD^33, 23^. In accordance with the effect of transcription-blocking DNA damage on gene-length-dependent gene expression^24^, we observed that in dauer samples longer genes are underexpressed after UV exposure. During dauer recovery, the animals efficiently recovered the overall gene expression programs including the GLTD during the course of their rejuvenation process. This suggests that during dauer exit DNA repair capacities are highly effective in restoring transcription integrity.

Altogether, we suggest a set of biological processes that slow aging during diapause and rejuvenate the organism, thus providing a model of naturally occurring longevity and rejuvenation, as reflected by functional assay and postdauer fitness, the BiT Age predictions, transcriptomic patterns in and during recovery from dauer, and recovery from transcription-blocking DNA damage.

## Acknowledgements

We thank the Regional Computing Center of the University of Cologne (RRZK) for providing computing time on the DFG-funded High Performance Computing (HPC) system CHEOPS as well as support. Nematode strains were provided by the National Bioresource Project (supported by The Ministry of Education, Culture, Sports, Science and Technology, Japan), the Caenorhabditis Genetics Center (funded by the NIH National Center for Research Resources, US), and the C. elegans Gene Knockout Project at the Oklahoma Medical Research Foundation (part of the International *C. elegans* Gene Knockout Consortium).

B.S. acknowledges funding from the European Research Council (ERC-2023-SyG, 101118919), the Hevolution Foundation (HF-GRO-23-1199212-35), the Deutsche Forschungsgemeinschaft (Reinhart Koselleck-Project 524088035, FOR 5504 project 496650118, FOR 5762 project 531902955, SFB 1678, SFB 1607, GRK 2407, CECAD EXC 2030 – 390661388, ANR-DFG project 545378328, and the DFG project grants 558166204, 540136447, 496914708, 437825591, 437407415, 418036758), the José Carreras Leukemia Foundation, DJCLS 04 R/2023, the Deutsche Krebshilfe (70114555), and the John Templeton Foundation Grant (61734).

## Methods

### Strain maintenance

Worms were kept in nematode growth medium (NGM) and fed with the *Escherichia coli* strain OP50. All strains were maintained at 15°C rather than the standard temperature of 20°C in order to avoid dauer formation in daf-c strains. For dauer induction and maintenance, worms were kept at 25°C until dauer exit. N2 Bristol strain was used as wild type (WT). The following strains were also used in this study:

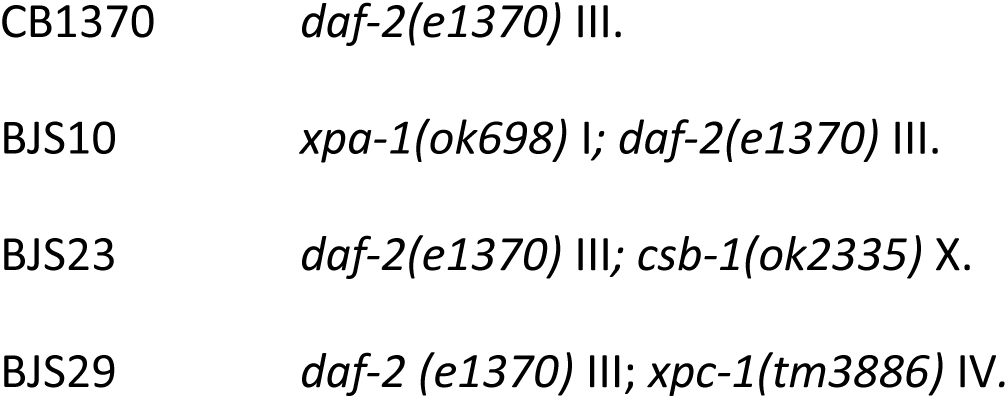

### Dauer induction and exit

Dauer entry was inducted by incubating bleach-synchronized L1 worms at 25°C on plates containing OP50. We considered day 1 dauers as worms incubated at 25°C for 72h, as they displayed dauer-specific anatomical characteristics, such as constricted pharynx and dauer alae, and resisted treatment with 1% sodium dodecyl sulfate (SDS) for 10 minutes. To induce exit from dauer, worms were washed off plates with 1x M9 buffer, transferred to plates freshly seeded with OP50, and incubated at 15°C.

### Bulky lesion-induction via UV exposure

For UVB broadband radiation treatment of worms, a Waldmann UV236B device equipped with Philips UV6 bulb (230V, 50Hz) was used. Lamp irradiance was measured prior to every irradiation using a UVX digital radiometer and a UVX-31 prove from UVP. Dosages used are specified in figure legends.

### Developmental resumption assay

Dauer exit was induced as described previously and developmental stages were assessed every 24h using a Zeiss Axio Imager Z1 microscope. Each plate analyzed contained about 150 to 200 worms.

### Brood size assay

Worms at the L4 stage were picked to plates seeded with OP50 and transferred to fresh plates every day until they ceased laying eggs. The number of viable progeny was assessed 48h after the parental worms were transferred to a fresh plate. For each plate analyzed, 15 to 20 L4 worms were initially picked. Worms that crawled off the plate before ceasing egg laying, or displayed internal hatching or ruptured vulva were excluded from analysis.

### Lifespan assay

Worms at the L4 stage were picked to plates seeded with OP50 and transferred to fresh plates every 48h until they ceased laying eggs. All lifespan assays performed in this study were done at 15°C. Scoring for alive worms was done every 24h during the initial 40 days and every 48h afterwards. Worms failing to show response to mechanical stimulation with a platinum wire or pharyngeal pumping were logged as dead. Worms that crawled off the plate before the end of the assay, or that displayed internal hatching or ruptured vulva were excluded from the analysis. For each replicate, 50 to 60 worms were initially picked for each condition.

### CPD repair assay via slot blot

Dauer populations were obtained by inducing dauer entry as described previously. Non-dauer groups were synchronized at the L3 stage about 72h after bleach-synchronized L1 populations were plated in plates seeded with OP50 and incubated at 15°C. Samples were recovered from plates using 1x M9 buffer and washed three times immediately after UV irradiation or after 24h or 72h. After washing, samples were quick-frozen using liquid nitrogen and stored at −80°C until DNA extraction was performed with Gentra Puregene Tissue Kit (Qiagen), according to the provided instructions. DNA concentration was measured with a Qubit device.

For each sample, 0.1 μg of DNA was used, with DNA hydration solution being added accordingly in order for all samples to have the final volume of 100 μl. Dilutions (1:2, 1:4, 1:8) were prepared and DNA was denatured for 5 min at 95°C using a S1000™ Thermal Cycler PCR machine (Bio-Rad). Samples were transferred to a Hybond nylon membrane (Amersham) using a Whatman 48-well slot blotting device (300 mbar vacuum). The membrane was incubated for 2h at 80°C for DNA cross-linking. The membrane was blocked in a 3% milk/TBS-T solution for 1 h at room temperature. Incubation with primary antibody (anti-CPD mouse anti-human from Cosmo Bio, 1:10000 in TBS-T) was done overnight at 4°C. Membrane was washed three times with TBS-T and blocked again in a 3% milk/TBS-T solution for 1 h at room temperature. Membrane was washed three times with TBS-T before being incubated with secondary antibody (peroxidase-conjugated AffiniPure goat anti-mouse, Jackson ImmunoResearch Laboratories, Inc., 1:10000 in TBS-T) 1h at room temperature. The membrane was washed three times with TBS-T and revealed in a dark room using Pierce™ ECL Plus Western Blotting Substrate (Thermo Fisher Scientific) and CL-XPosure™ Film (13×18 cm, Thermo Fisher Scientific). To assess total DNA levels, the membrane was washed five times with TBS-T and incubated with SYBR Gold solution (Thermo Fisher Scientific, 1:10000 in TBS-T) overnight at 4°C. Membrane was revealed using a ChemiDoc Imaging System (Bio-Rad) and the Bio-Rad ImageLab 5.0 software.

### RNA purification and sequencing

Each condition contained three biological replicates of 8000 worms, collected in ice-cold 1x M9 buffer from plates seeded with OP50. The samples were washed three times with 1x M9 buffer, resuspended in Trizol (Invitrogen) and 0.7 mm zirconia beads (Roth), and quick-frozen in liquid nitrogen at −80°C until RNA extraction was performed. The samples were disrupted using a Precellys24 homogenizer (Bertin) and 1-Bromo-3-chloropane (Sigma-Aldrich) was added for phase separation. The purification of RNA was performed using the RNeasy Mini Kit (Qiagen) according to the provided instructions. The isolated RNA was eluted in RNase-free H2O and quantified using a NanoDrop 8000 spectrophotometer (Thermo Fisher Scientific). All steps of quality control and sequencing of RNA were performed at the Cologne Center for Genomics (CCG) using an Illumina Hiseq 4000 device. The sequencing was paired end and produced approximately 15 million reads per sample.

The quality of produced raw sequencing files was assessed using FastQC reports supplied by CCG. The raw files were pre-processed using fastp^34^ that performs read filtering, adapter trimming, etc. The read quantification was performed using Salmon^35^ with –gcBias, -l A and -- validateMappings. The index is based on WBPS15 reference mRNA transcriptome with the corresponding reference genome used as decoy for the indexed reference with -k 31. The data can be retrieved from gene expression omnibus under GSE288723.

### Statistical analysis and graphs

All statistical methods used, number of biological replicates and error bar descriptions are provided in the respective figure legends or graphs. Sample size is indicated in each figure or respective method section. Data and statistical significances were analyzed with the GraphPad Prism 7 software package, unless otherwise specified. Significance levels are ∗p<0.05, ∗∗p<0.01 and ∗∗∗p<0.001. Worms were always randomly selected from large populations for each experiment performed.

### BiT Age analysis

Biological age prediction was performed using the BiT Age clock as previously described^13^. Briefly, gene expression data were binarized such that genes with expression levels above the median expression value (after removing zero-count genes) were set to 1, while all others were set to 0. The BiT Age score for each sample was then calculated by summing the predefined coefficients of the clock genes that were assigned a 1 after binarization, followed by the addition of the BiT Age intercept to obtain the final predicted biological age. We used the updated BiT Age version 2, which is available at https://github.com/Meyer-DH/AgingClock

### Gene expression analysis

tximport^36^ was used together with caenorhabditis_elegans.PRJNA13758.WBPS15.canonical_geneset.gtf to summarize the transcript expressions obtained via salmon on a gene level. We used DESeq2^37^ to create a generalized linear model (GLM) for each gene, one for dauer aging and the other for dauer exit, with dauer day and exit hour as the continuous design variable, meaning the slope of the model is the log2 fold change (LFC) of the gene for one unit change. Implicated pathways were investigated by conducting KEGG enrichment analyses. In order to identify genes whose dauer exit expression dynamic changes with dauer duration, a linear model was constructed for each gene with dauer day and exit hour. For every differential expression analysis, a pre-filtering step was implemented to retain only genes with counts of at least 10 in at least 3 samples, which is the smallest group size. For the creation of the PCA, the samples were also variance stabilized via vst. Time series differential expression analysis was carried out with the parameters test = “LRT” (Likelihood ratio test) and a reduced model excluding the respective factor of interest.

For the dauer aging gene clustering, the differentially expressed genes were per-gene normalized and k-means clustered into 4 groups, as determined via the elbow criterion, using the kmeans function with iter.max = 1000 and nstart = 100.

enrichKEGG^38^ was used for pathway enrichment with organism = ‘cel’, keyType = ‘ncbi-geneid’, pAdjustMethod = “BH” and pvalueCutoff = 0.05. The wormbase ids of the genes were converted to entrez ids via mapIds with multiVals = “asNA” using org.Ce.eg.db as the AnnotationDb object. The universe was set to all genes in the dataset that passed DESeq2 independent filtering. Only pathways with an adjusted p-value of at most 0.05 were considered. The analyses were carried out on the 18.03.2025.

The dauer exit signatures were created by selecting genes that are differentially expressed as a function of exit hour on the respective day. Genes with a positive LFC were considered to be upregulated and genes with a negative LFC were considered to be downregulated.

For the gene length comparison, the start and end positions of the respective genes were retrieved from biomaRt^39^.

Differential gene expression in response to UV exposure was assessed using three pairwise comparisons: D4 UV vs. D4, D4 UV 6h vs. D4 6h, and D4 UV 24h vs. D4 24h. Genes with a false discovery rate (FDR) below 0.05 in at least one comparison were retained for visualization. Log₂ fold change (logFC) values were used as input for clustering. Genes were grouped into four clusters using k-means clustering, based on their expression patterns across the dauer aging time course (days 1, 4, 15, and 30). Columns (comparisons) were hierarchically clustered using the Ward method with Euclidean distance. Heatmaps were generated using the clustermap function from the Seaborn Python library.

## Supplementary figure legends

**Supplementary Figure 1.** Shown are all significantly enriched KEGG pathways for genes exhibiting differential expression across the dauer aging time course. Color-coded is the adjusted p-value, the bubble size shows the rich factor. This figure expands on the subset of pathways presented in **Figure 3A**.

**Supplementary Figure 2.** Shown are all significantly enriched KEGG pathways for each of the four gene clusters identified across the dauer aging time course. The x-axis represents the gene clusters (Groups 1–4), while the y-axis lists all enriched pathways. Color-coded is the adjusted p-value, the bubble size shows the rich factor. This figure expands on the subset of pathways presented in **Figure 3C**.

**Supplementary Figure 3.** Dauer exit is similar between D1 and D4 UV-treated dauers. *daf-2* animals were UV- or mock-treated at day 1 of dauer. Panels (A) to (D) show, respectively, 0, 15, 30 and 45 mJ/cm2 conditions. Dauer exit was induced immediately after UV treatment. (average of n=3 independent experiments per dose is shown, error bars represent the standard deviation (SD). Two-tailed t-test. No significant differences detected between D1 and D4).

**Supplementary Figure 4.** (A) Box plots of gene lengths of genes differentially over- or under-expressed in UV-treated samples of dauer exit on D4 relative to untreated samples of dauer exit on D4 with n being the number of genes in the set. Mann Whitney U test was used to identify statistically significant differences in gene lengths between the groups. (p< 0.05 *, p < 0.01 **, p < 0.001 ***). (B) Heatmap showing the log₂ fold change (logFC) of genes that are significantly differentially expressed (FDR < 0.05) in at least one of the three comparisons: D4 UV vs. D4, D4 UV 6h vs. D4 6h, and D4 UV 24h vs. D4 24h. Rows represent individual genes, and columns represent the pairwise comparisons. Genes were clustered using hierarchical clustering with the Ward method and Euclidean distance. The color scale indicates the direction and magnitude of differential expression (blue: downregulated; red: upregulated).

## Supplementary Table legends

**Supplementary Table 1.** Source data and corresponding statistical analyses of Figure 1.

**Supplementary Table 2.** Source data and corresponding statistical analyses of the BiTAge predictions (Figure 2C).

**Supplementary Table 3.** KEGG enrichment results of the dauer aging (Fig. 3A), cluster aging (Fig. 3C).

**Supplementary Table 4.** KEGG enrichment results of dauer exit (Fig. 4B)

**Supplementary Table 5.** Source data and corresponding statistical analyses of Figure 5.

**Supplementary Table 6.** Differentially expressed genes and their lengths (Fig. 6).

**Supplementary Table 7.** Source data for dauer exit comparison data under UV (Suppl. Fig. 3).

