## Supplementary figures and images for "Age deceleration and reversal gene patterns in dauer diapause"

### Supplemental Figure 1

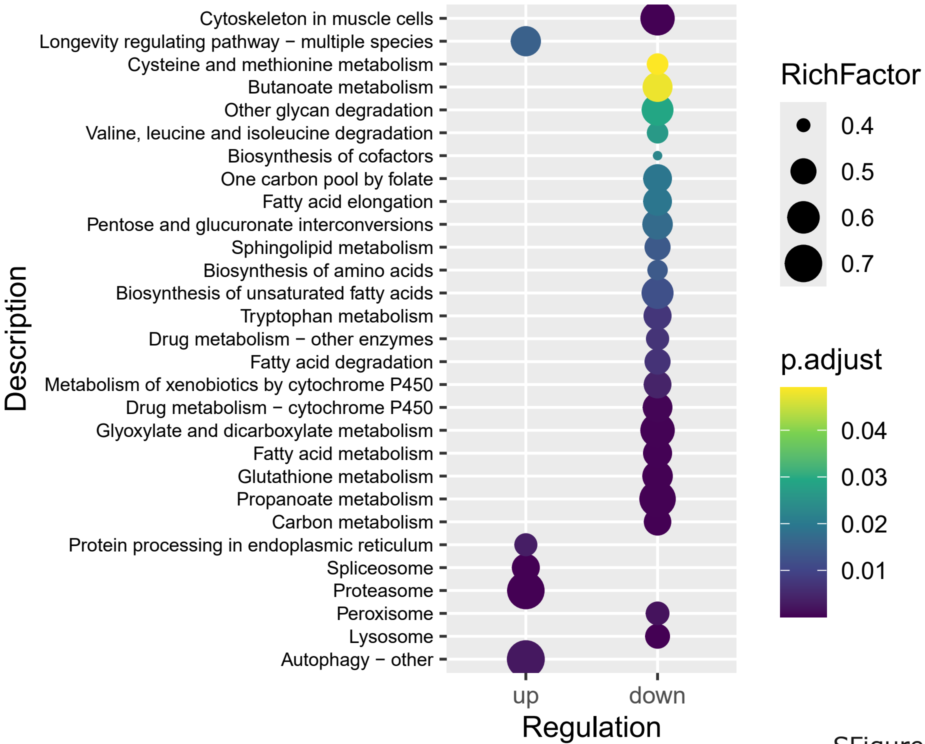

### Supplemental Figure 2

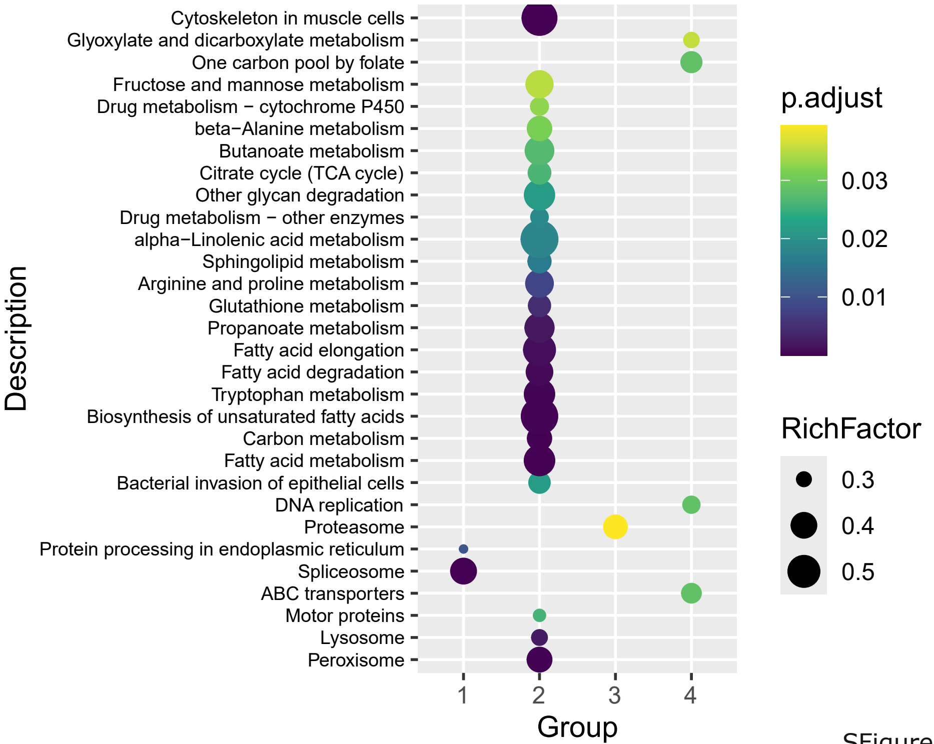

### Supplemental Figure 3

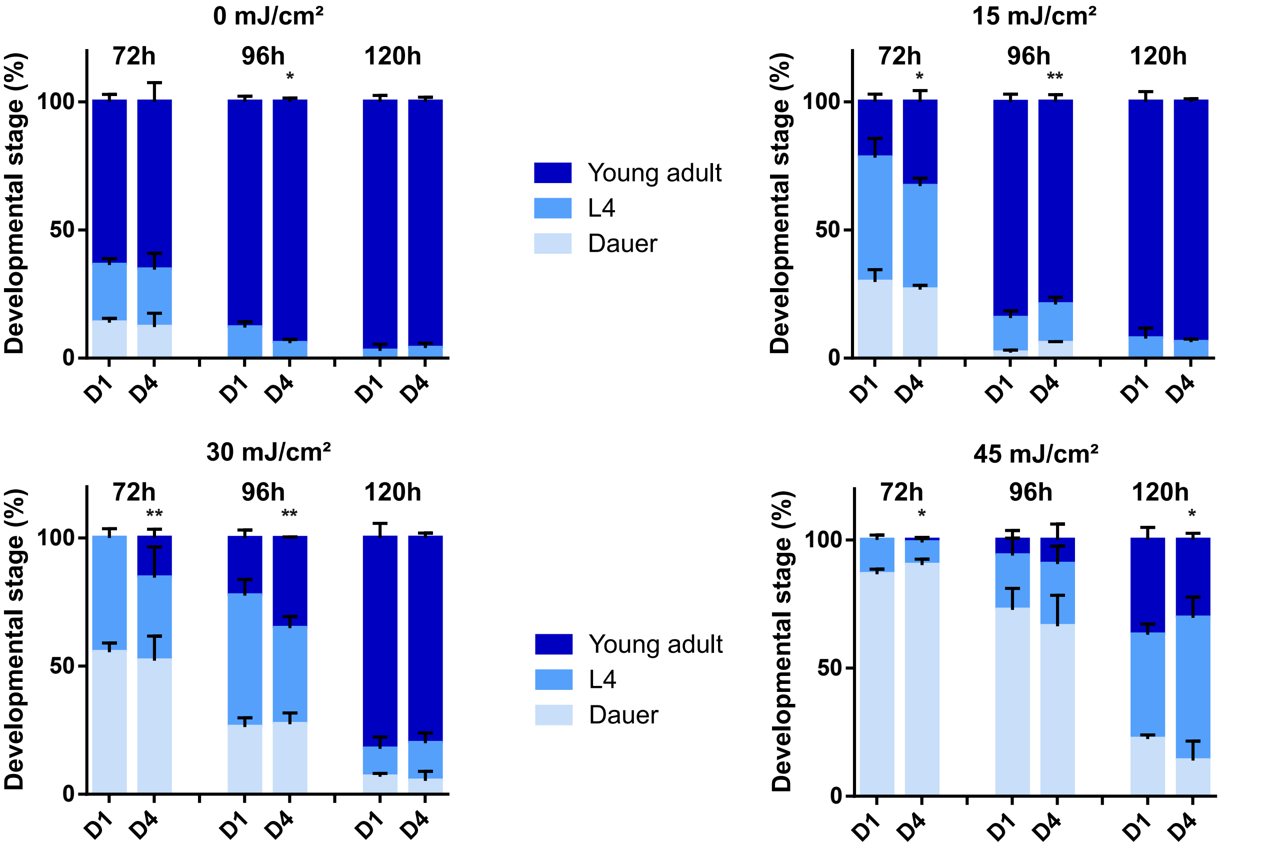

### Supplemental Figure 4

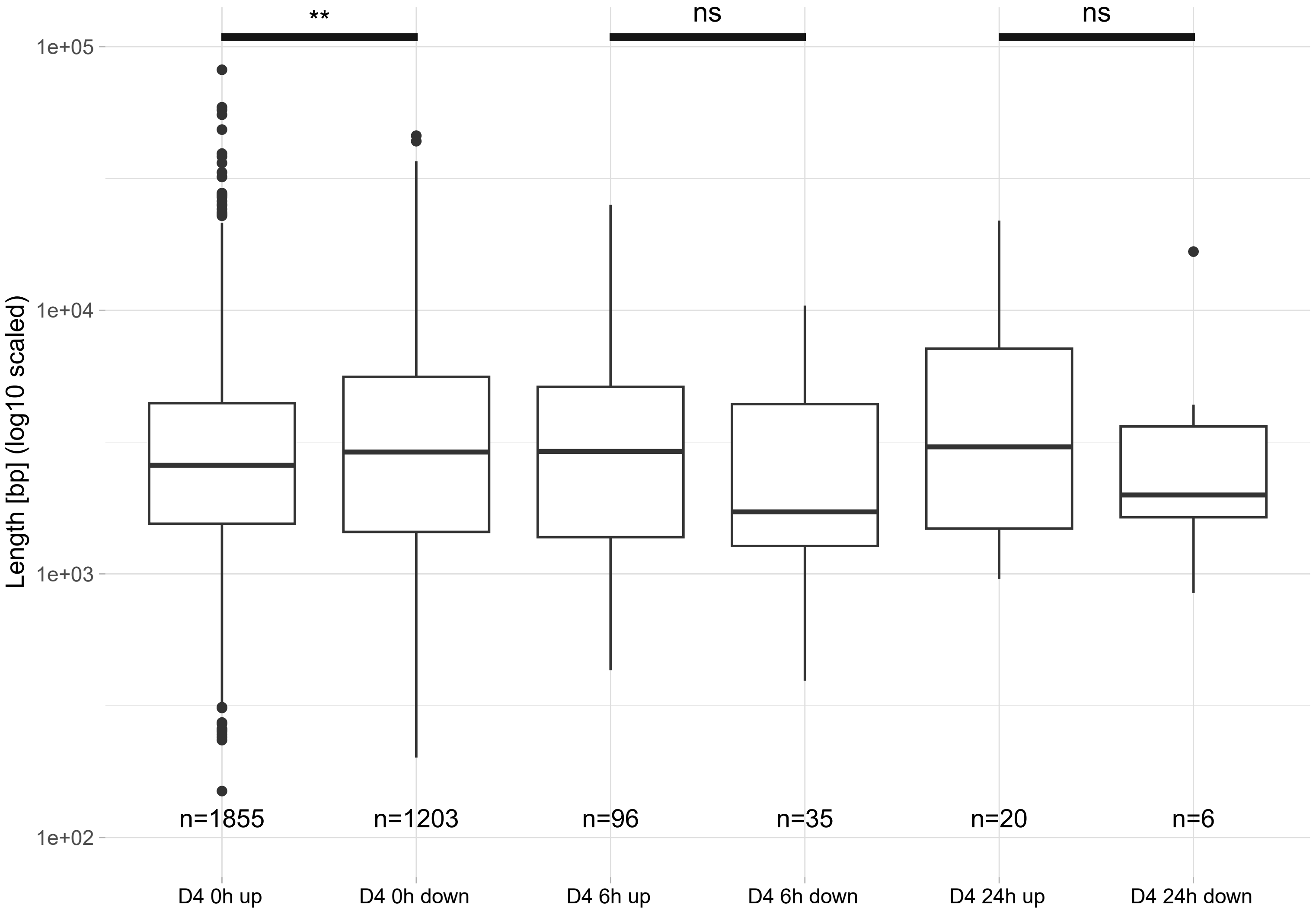

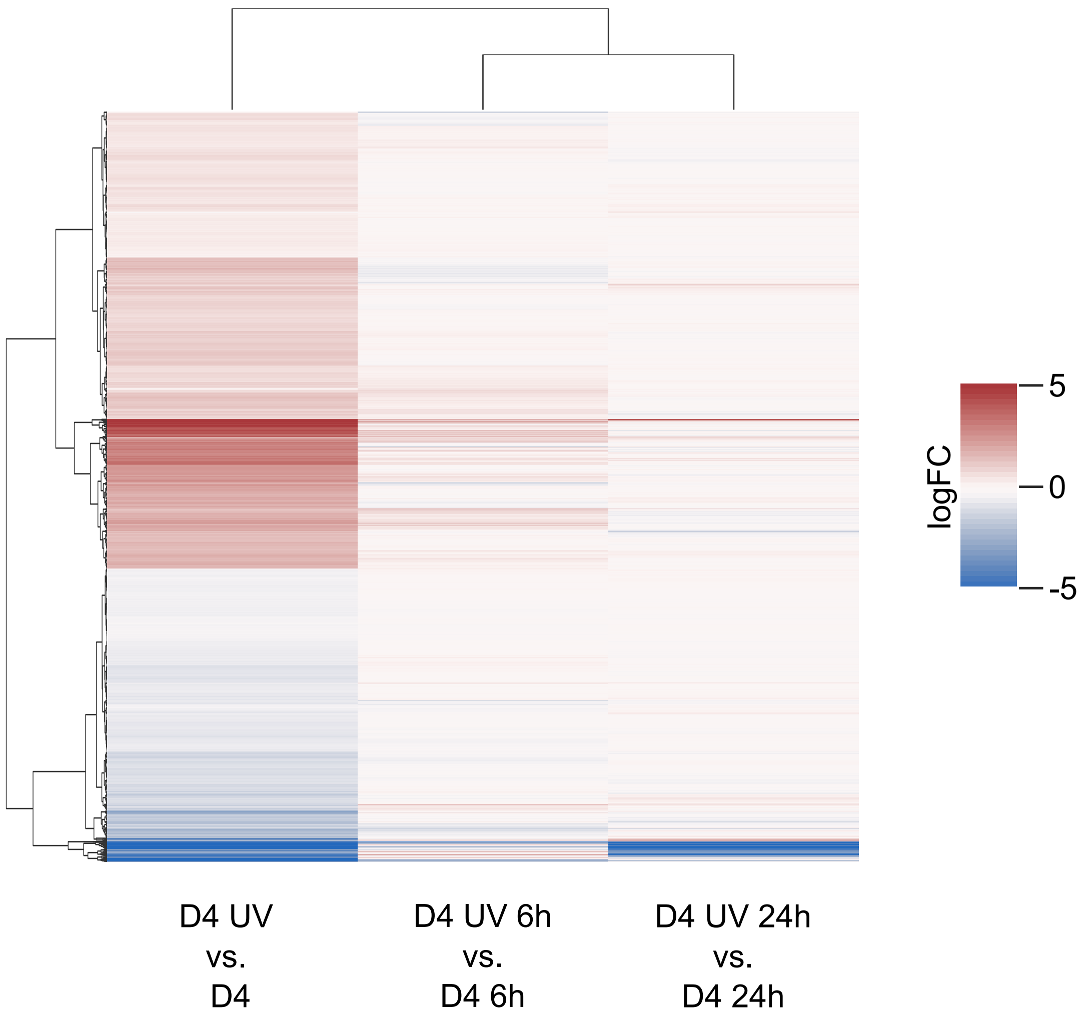
